# A simple and cost-effective method for generating spheroids from triple-negative breast cancer cell line (MDA-MB-231)

**DOI:** 10.64898/2025.12.17.694917

**Authors:** Ramón Cervantes-Rivera, Luisa Nirvana González-Fernández, Atalia Ziret Romero Rosas, Sandra Jetsamari Figueroa Ortíz, Alejandra Ochoa Zarzosa, Joel E. López-Meza

**Affiliations:** Centro Multidisciplinario de Estudios en Biotecnología (CMEB), Facultad de Medicina Veterinaria y Zootecnia, Universidad Michoacana de San Nicolás de Hidalgo, Morelia, Michoacán, México; Secretaría de Ciencia, Humanidades, Tecnología e Innovación

**Keywords:** Triple-negative breast cancer, 3D cell culture, Spheroids, Drug screening, Tumor microenvironment

## Abstract

Breast cancer (BC) is the most frequently diagnosed malignancy in women and a leading cause of cancer-related mortality worldwide. Molecular classification based on estrogen receptor (ER), progesterone receptor (PR), HER2, and Ki67 expression guides prognosis and therapy, with triple-negative breast cancer (TNBC)—lacking ER, PR, and HER2 representing 15–20% of cases. TNBC’s aggressive behavior, early recurrence, and limited treatment options underscore the need for improved models to develop targeted therapies. While monolayer (2D) cultures have advanced cancer research, they poorly replicate the three-dimensional (3D) tumor microenvironment (TME), leading to translational gaps. 3D spheroids address these limitations by recapitulating cell-cell/matrix interactions, metabolic gradients, and hypoxic cores, offering a physiologically relevant platform for studying metastasis, drug resistance, and therapeutic screening. Here, we present a simple, cost-effective method for generating spheroids. This protocol is applicable across different cell types, bridging the gap between traditional 2D cultures and *in vivo* studies. 3D cell culture opens the door to personalized medicine and drug discovery.

**Graphical overview:** 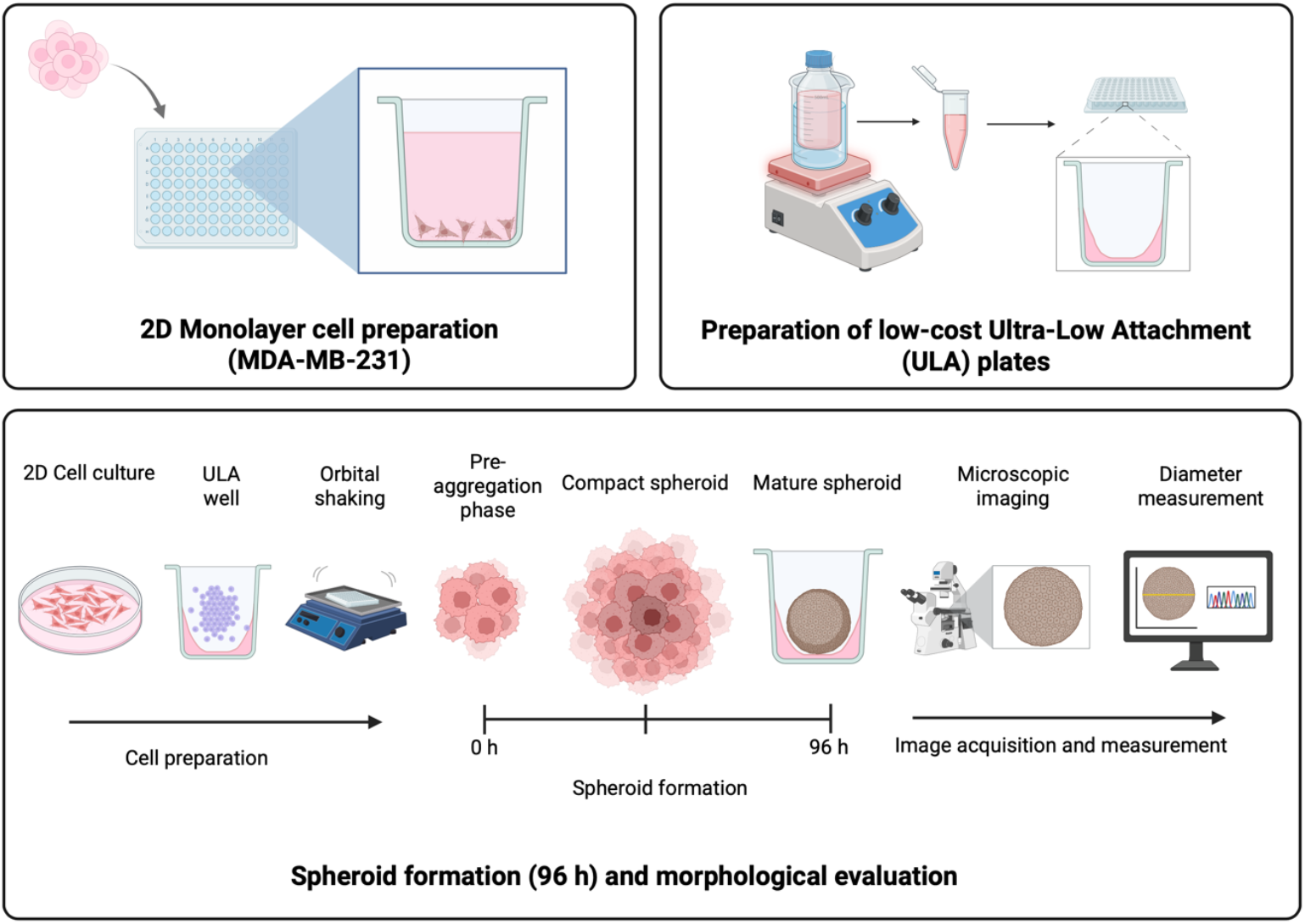

**Key features:**

- Employs a cost-effective, lab-made agarose coating to create ultra-low attachment surfaces in standard 96-well plates.
- Specifically optimized for generating consistent spheroids from the aggressive MDA-MB-231 triple-negative breast cancer cell line.
- Generates measurable spheroids within 96 hours using only basic cell culture equipment and an orbital shaker.
- Provides a clear workflow from spheroid formation to quantitative size analysis using freely available (ImageJ) and common (GraphPad Prism) software.

## Background

Breast cancer (BC) remains the most diagnosed malignancy in women globally and the leading cause of cancer-related deaths [1]. BC is classified into molecular subtypes based on the expression of estrogen receptor (ER), progesterone receptor (PR), human epidermal growth factor receptor 2 (HER2), and the proliferation marker Ki67 [2, 3]. These subtypes dictate prognosis and guide therapeutic strategies, with hormone receptor-positive (ER+/PR+) and HER2-enriched tumors benefiting from targeted therapies. Among these, triple-negative breast cancer (TNBC)—defined by the absence of ER, PR, and HER2 expression—represents 15–20 % of all breast cancers and is associated with aggressive clinical behavior, early recurrence, and limited treatment options [4]. Due to its lack of targetable receptors, TNBC patients primarily rely on chemotherapy, highlighting the urgent need for novel therapeutic approaches.

For decades, monolayer (2D) cell cultures have been the cornerstone of cancer research, enabling foundational discoveries in cell proliferation, migration, and drug screening [5]. However, these models fail to recapitulate the three-dimensional (3D) architecture and tumor microenvironment (TME) of *in vivo* tumors, leading to discrepancies in drug responses and mechanistic studies [6]. To bridge this gap, 3D spheroid models have emerged as a powerful tool, better mimicking the complexity of solid tumors [7]. These models incorporate cell-cell and cell-matrix interactions, polarization and spatial organization, oxygen and nutrient gradients, metabolic heterogeneity, and hypoxic cores that resemble *in vivo* conditions. As a result, spheroids provide a more physiologically relevant platform for studying tumor invasion and metastasis, drug resistance mechanisms, therapeutic screening, and personalized medicine approaches [8, 9].

Recent advances in scaffold-based and scaffold-free spheroid generation techniques—such as hanging drop, ultra-low attachment plates, and microfluidic systems—have further improved reproducibility and scalability [10, 11]. The spheroid generation method presented here offers a simple, cost-effective, and universally applicable strategy to harness the power of 3D cell culture. Its ability to produce reliable *in vitro* models that more accurately mimic the *in vivo* conditions is a crucial step forward. We anticipate that the widespread adoption of this protocol will not only enhance the predictive power of preclinical drug screening but also serve as a cornerstone in the foundational research required to usher in a new era of personalized therapeutic interventions.

## Materials and reagents

1. MDA-MB-231 cell line (ATCC, catalog number: CRM-HTB-26)
2. Dulbecco’s modified Eagle medium (DMEM) (Sigma Aldrich, catalog number: A2942)
3. L-glutamin (Gibco, catalog number: 20530-081)
4. Amphotericin (SIGMA, catalog number: A2942)
5. Fetal bovine serum (FBS, BioWest catalog number: BIO-S1400)
6. Bovine calf serum (BCS, BioWest catalog number: S0400-500)
7. Penicillin-streptomycin (GIBCO, catalog number: 15120122)
8. Trypsin (SIGMA, catalog number: T4799-5G9) (see Recipes)
9. Ethylenediaminetetraacetic acid (EDTA) (J.T Baker, catalog number: 8993-01)
10. Phosphate-buffered saline (PBS) (see Recipes):
  a. Sodium chloride (J.T Baker, catalog number: 3624-01)
  b. Sodium phosphate, 7-hydrate, crystal (J.T Baker, catalog number: 3824-01)
  c. Monobasic potassium phosphate crystal (J.T Baker, catalog number: 3824-01)
  d. Potassium chloride (J.T Baker, catalog number: 3040-01)
11. Ultra-low attachment 96-well plates (see Recipes)
12. Pipette tips
  a. 1,000 μL (Uniparts, catalog number: 51131)
  b. 200 μL (Uniparts, catalog number: 51121Y)
  c. 10 μL (Axygen, catalog number: T-300)
13. Agarose (Invitrogen, catalog number: 16500-100)
14. MQ-water sterile water
15. Nitrile gloves
16. Culture tube of 15 mL (Uniparts, catalog number: 34117F)
17. Cell culture dish, 60 mm X 15 mm (Nest, catalog number: 705001)
18. Cell culture plate, 96 wells, flat bottom, TC (Nest, catalog number: 701001)

### Recipies

#### a) Culture medium for MDA-MB-231 cells Dulbecco’s modified Eagle medium (DMEM)

1. Dissolve DMEM powder in sterile distilled water and stir until completely dissolved.
2. Add 1.5 g sodium bicarbonate (NaHCO_3_).
3. Adjust the pH to 7.4 by carefully adding drops of NaOH (1 M) or HCl (1 M), as needed.

Under a laminar flow hood, supplement the medium with the following components:

1. 5% FBS (25 mL)
2. 5% ST (25 mL)
3. 1% Penicillin–streptomycin (5 mL)
4. 0.05% Amphotericin B (250 µL)
5. 0.05% L-Glutamine 200 mM (250 µL) Mix gently to homogenize and store at 4 °C until use.

#### b) Agarose 1.5 %

1. Add 1.5 g agarose to 100 mL triple-distilled water in a glass flask.
2. Sterilize the solution by autoclaving for 15 minutes.

#### c) PBS 1X

1. Autoclave a glass flask containing 1 L of triple-distilled water and allow it to cool down at room temperature.
2. Add the following salts to the flask: 8 g NaCl, 0.2 g KCl, 1.44g Na_2_HPO_4_, and 0.24 g KH_2_PO_4_.
3. Dissolve the salts using a magnetic stirrer.
4. Calibrate the pH meter and adjust the solution to pH 7.4 by slowly adding 1 M NaOH or 1 M HCl, as needed.
5. Remove the stir bar, filter the solution through a 0.22 µm membrane under sterile conditions, and store at 4 °C until use.

#### d) Trypsin-EDTA

1. Start with 1 L of PBS 1X.
2. Dissolve 390 mg trypsin and 85 mg NaCl in 10 mL of PBS 1X.
3. Add 370 mg EDTA to the remaining PBS and mix until dissolved.
4. Combine with the trypsin–NaCl solution and stir until fully mixed.
5. Filter the solution through a 0.22 µm unit under sterile conditions.
6. Label and store at –20 °C until use.

### Equipment

1. Centrifuge (PowerSpin™, model: C856)
2. Micropipettes (Axygen, catalog number):
  a. **AP-10**: 0.5–10 µL Single-channel Pipettor (Axygen® Axypet®)
  b. **AP-100**: 10–100 µL Single-channel Pipettor (Axygen® Axypet®)
  c. **AP-1000**: 100–1,000 µL Single-channel Pipettor (Axygen® Axypet®)
3. Orbital shaker (Benchmark, model: BT302)
4. Digital stirring hot plate (Thermo Scientific, model: SP131015Q)
5. Inverted microscope (Carl Zeiss, model: 37081)
6. Laminar flow cabinet (Thermo Fisher Scientific, model: 1340)
7. Incubator for cell culture (Thermo Fisher Scientific, model: 3422)
8. Neubauer chamber (Mariendfeld, catalog number: 0610010)
9. Microwave oven

### Software and datasets

1. ImageJ (https://imagej.net/ij/download.html)
2. GraphPad Prism 9 (https://www.graphpad.com/features)

### Procedure

Before starting to work with cell culture, it is crucial to clean all surfaces and equipment that will be used during the experimental procedure.

#### A) Cost-effective Ultra Low Attachment Plate preparation

1. Maintain the sterile 1.5% (w/v) agarose solution in a water bath set to 100 °C on a stirring hot plate.
2. Use a 96-well cell culture plate with a flat bottom and TC to prepare an Ultra Low Attachment Plate.
3. Using sterile technique, transfer 1 mL of the 1.5% molten agarose solution (see Recipes) into a 1.5 mL microcentrifuge tube.
4. Dispense 60 µL of the agarose solution into each well of the plate from the stock tube (Figure 1, 2).
5. Repeat the previous step until all wells are filled (Figure 2).
6. Allow the agarose to solidify, then uncover it at room temperature under a cell culture hood for 15 minutes.
7. Store the prepared plates in a sterile environment until their use.

**Figure 1.**
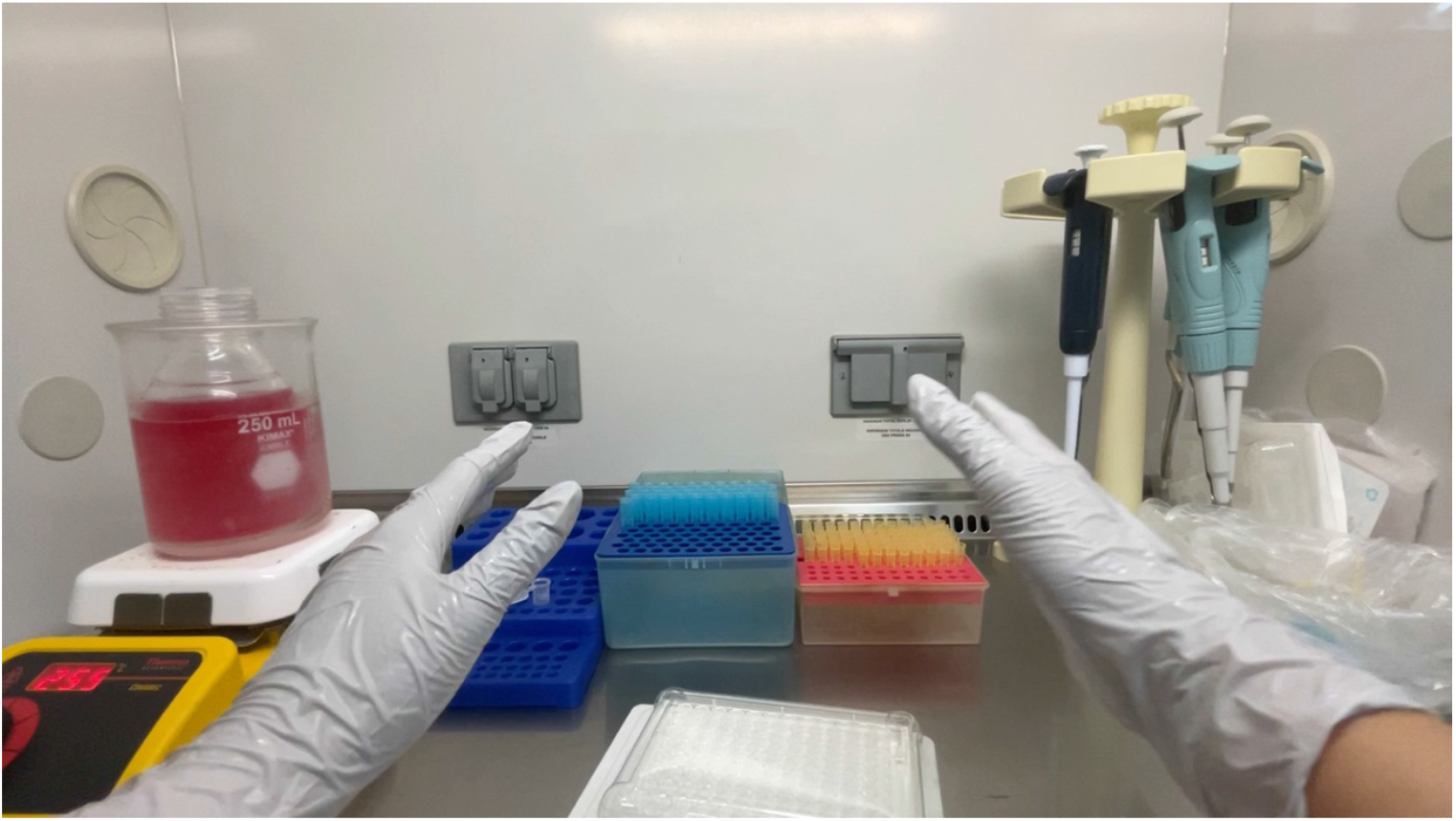
Cost-effective Ultra Low Attachment Plates production.

**Figure 2.**
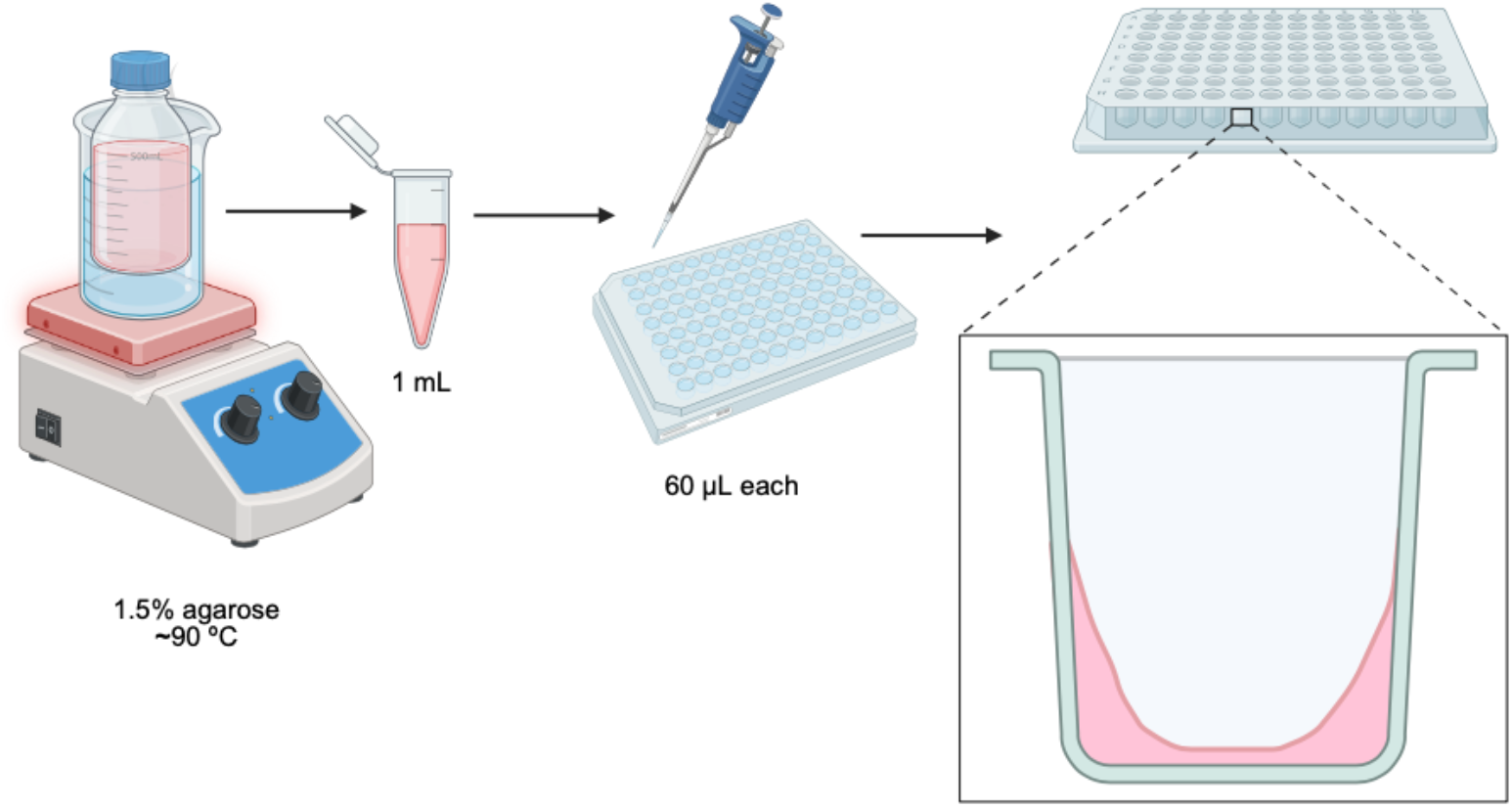
Schematic representation of ULA’s production.

#### B) Thaw MDA-MB-231 cells

1. Remove the vial containing the cells from the liquid nitrogen tank.
2. Place it in the incubator at 37 °C until the cells are completely thawed.
3. Transfer the unfrozen cells with the micropipette into a 15 mL culture tube.
4. Centrifuge the 15 mL tube containing cells for 5 minutes at 1,500 rpm at room temperature.
5. Remove the supernatant with a 1000 µL pipette.
6. Resuspend cells in 1 mL of DMEM complete.
7. Place them in a 60 mm x 15 mm cell culture dish and add 5 mL of complete DMEM.

#### C) Preparation of MDA-MB-231 cells

Grow MDA-MB-231 cells in a cell culture dish (60 mm X 15 mm) with complete medium (DMEM) at 37 °C, 5 % CO_2_ until ∼80% confluency.

1. Aspirate the medium and wash with 1X PBS.
2. Add 3 mL of trypsin-EDTA 0.25% and incubate at 37 °C for 3-5 min.
3. Neutralize trypsin with 3 mL of DMEM-complete medium.
4. Centrifuge at 1500 RPM for 5 minutes at room temperature. Discard the supernatant and resuspend the pellet in 1 mL of fresh DMEM-complete medium.

#### D) Count cells in the Neubauer chamber

1. Prepare a 1:10 dilution of cells treated with trypsin.
2. To prepare dilution 1:10, add 90 µL of DMEM-complete medium in a microtube (1.5 mL), and 10 µL of cells previously resuspended in DMEM-complete medium.
3. Take 10 µL of the diluted cells and place them in the Neubauer chamber. Count the cells and calculate the total number of cells (Figure 3).
4. Cell counting formula:

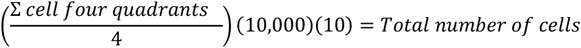

**Figure 3.**
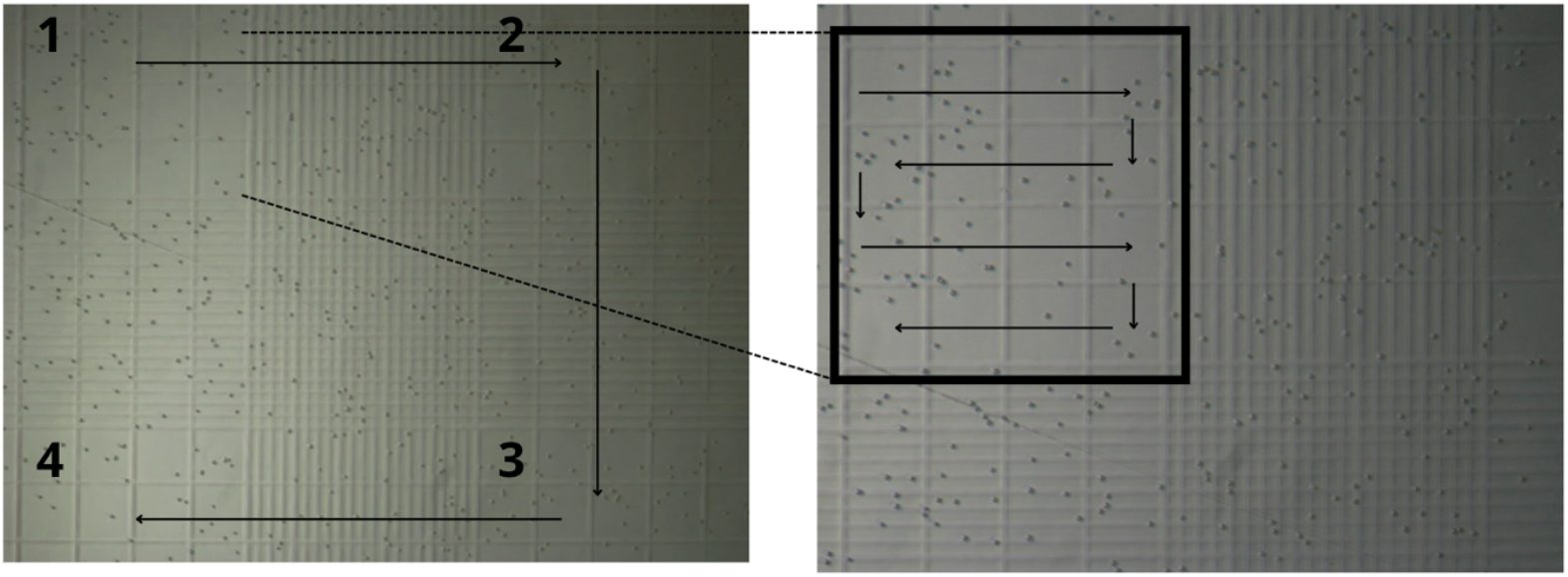
Representation of the counting cells on a Neubauer chamber.

#### E) Prepare dilutions with different concentrations of cells

To determine the optimal cell number for spheroid formation, it is essential to prepare different cell dilutions and place them in an Ultra Low Adherent Plate.

1. Prepare dilutions of 3 X 10^3^, 5 X 10^3^, 8 X 10^3^, 10 X 10^3^, 15 X 10^3^, 20 X 10^3^, 25 X 10^3,^ and 30 X 10^3^ (Figure 4).
2. In an Ultra Low Attachment 96-well plate, put 100 μL of each dilution.
3. Shake the plate in an orbital shaker for 3 h at 1500 RPM at room temperature.
4. After the shaker period, put the plate in the incubator at 37°C, 5% CO2, and 95% humidity.
5. Monitor the spheroid formation daily for 96 h (4 days).
6. Take a picture every 24 h in an inverted microscope with a 10X objective (Figure 5).

**Figure 4.**
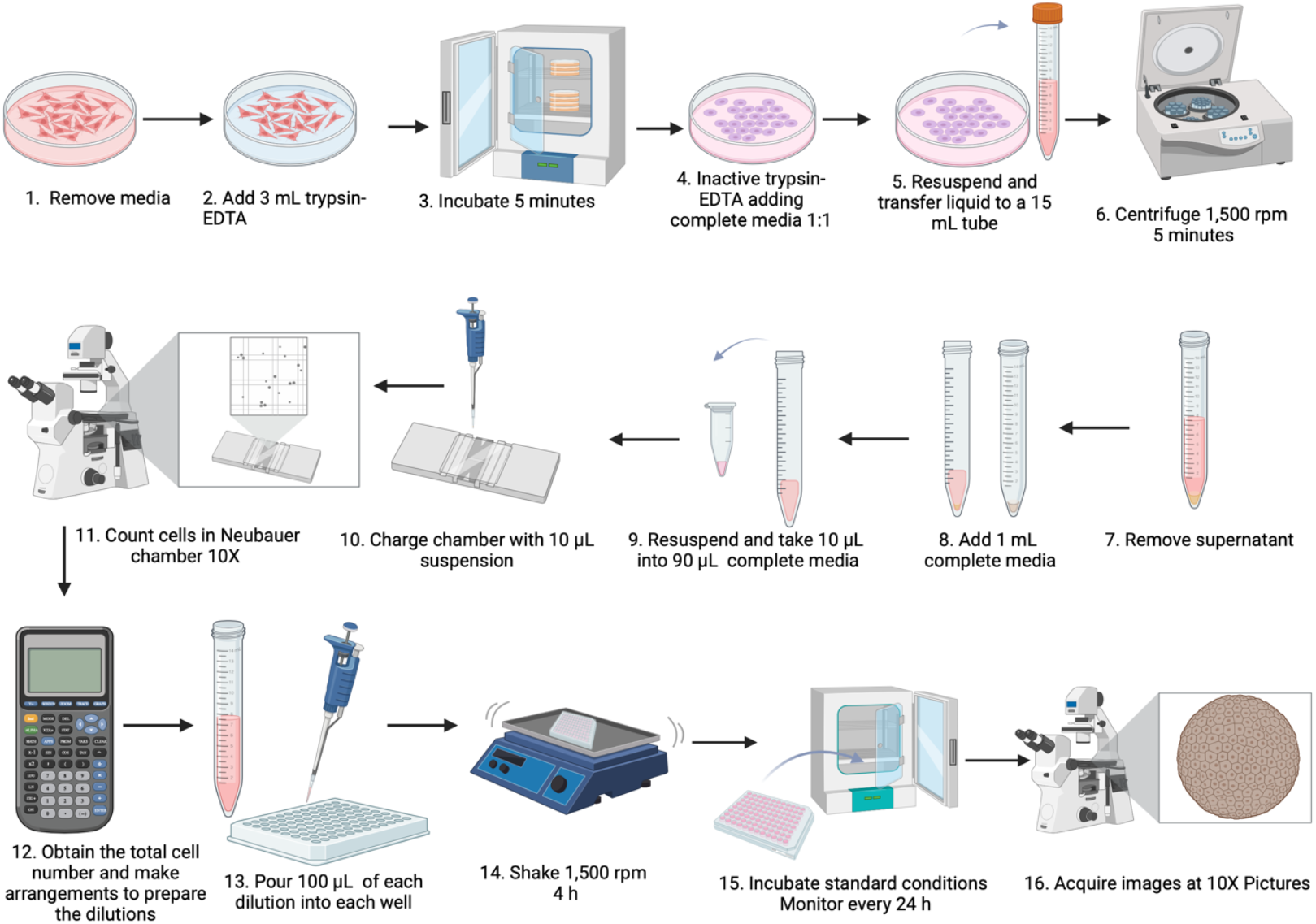
Schematic representation of low-cost spheroid formation method.

**Figure 5.**
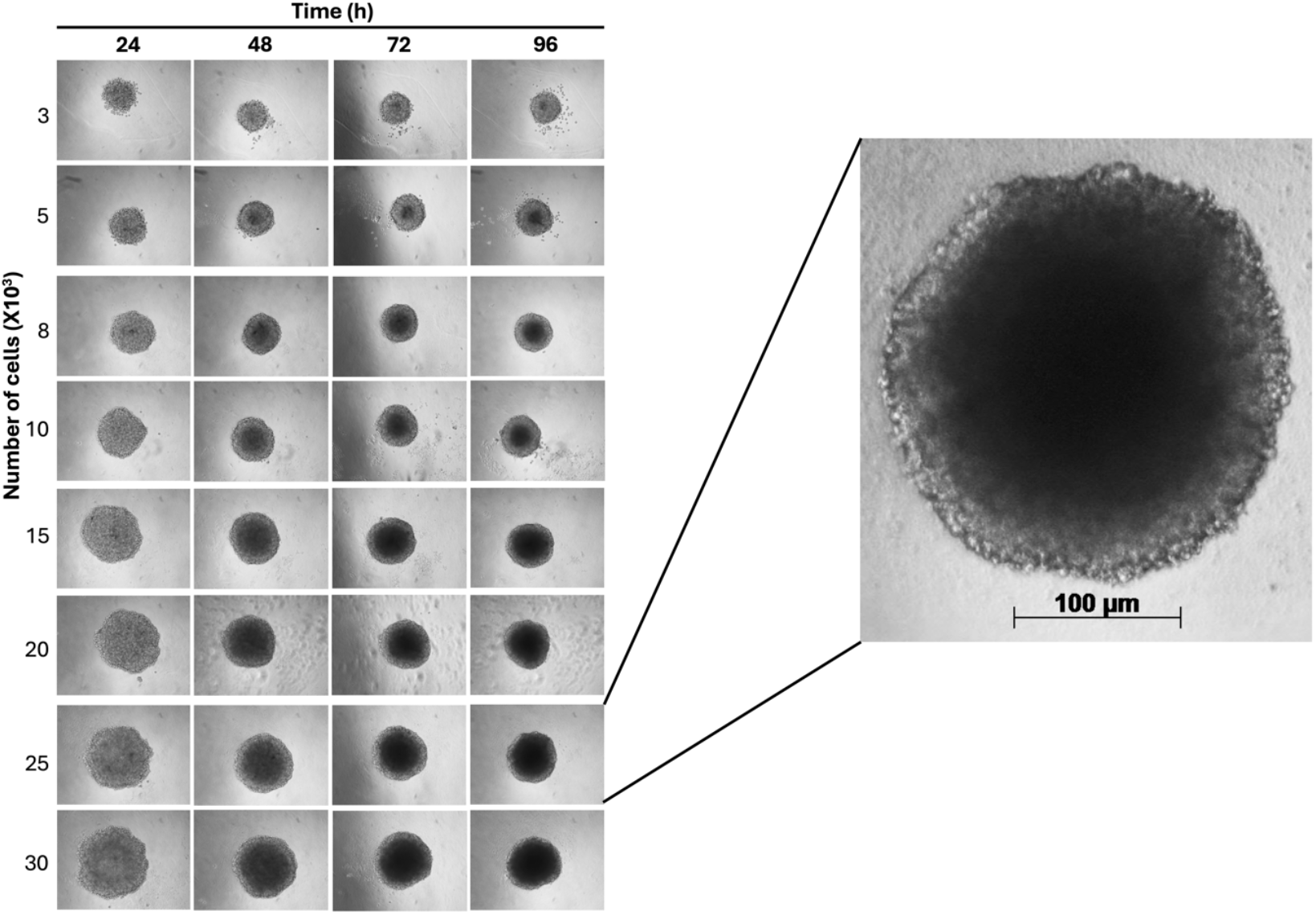
Step-by-step process for setting the scale and measuring spheroid circumference using ImageJ.

#### F) Monitoring and Analysis of Spheroids

1. Acquire images of all spheroids every 24 h using the inverted microscope (Carl Zeiss, 10X magnification).
2. Record a scale reference (micrometer or scale bar) on the microscope during image acquisition.
3. After 96 h of culture, transfer all images to *ImageJ* (NIH, Bethesda, MD, USA).
4. Set the global scale in *ImageJ* using the reference obtained from the microscope.
5. Measure the circumference of each spheroid in all images (Figure 6).
6. Export the measurement data and use them for analysis in *GraphPad Prism 9* (Figure 6 and 7).

**Figure 6.**
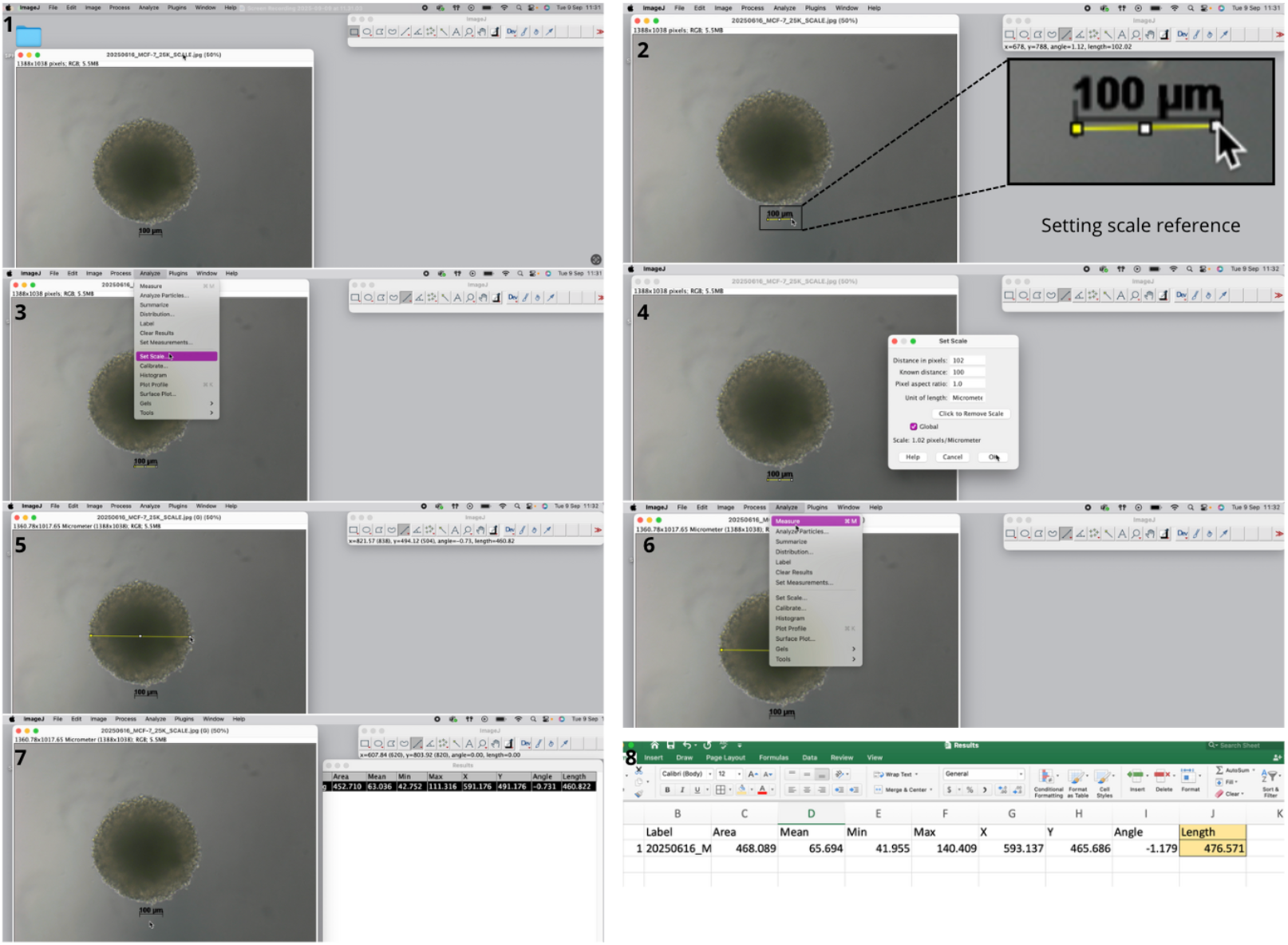
Representative imagens of spheroids along 96 h (10X magnification).

**Figure 7.**
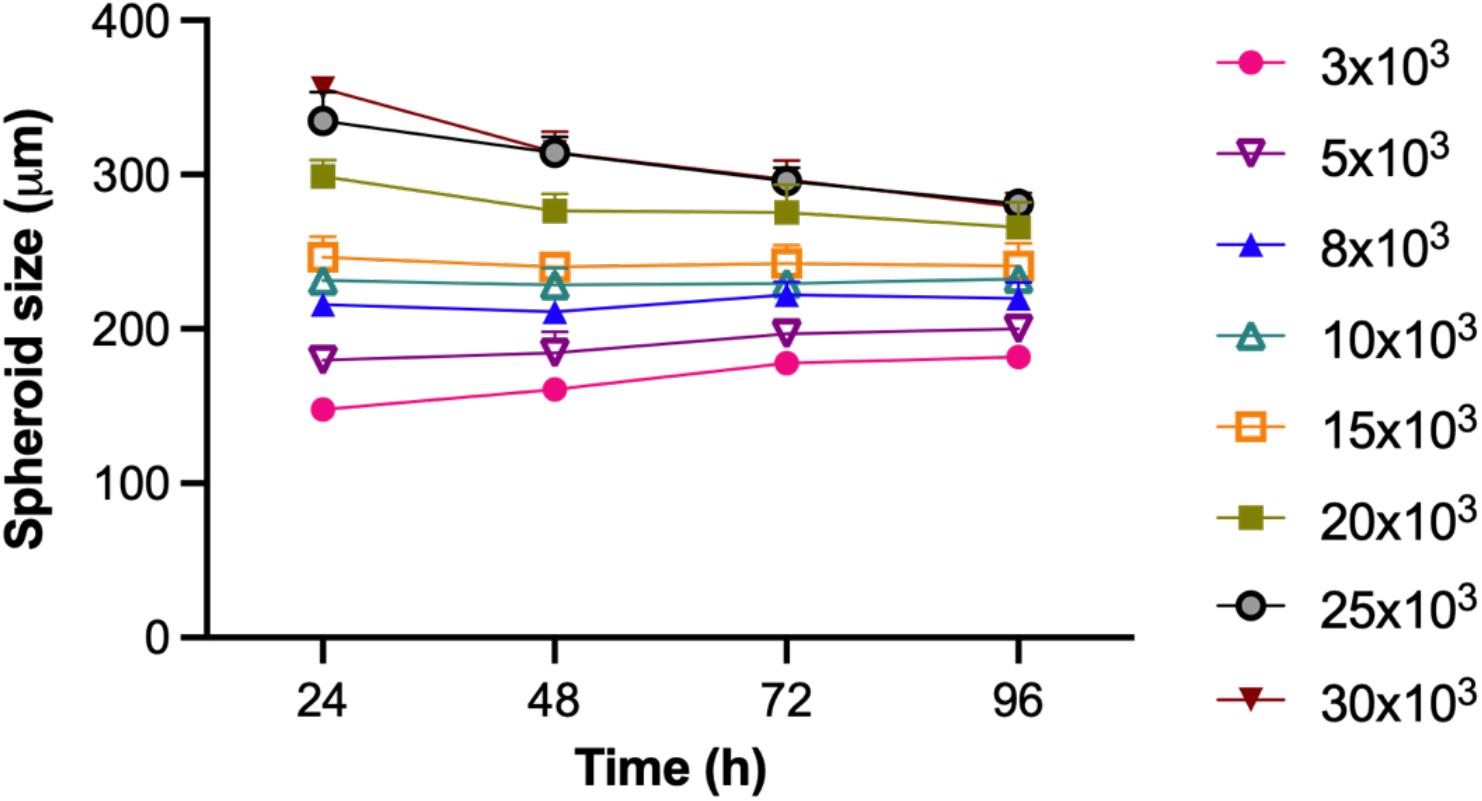
Evolution of spheroid size during the 96-h culture period. Each point represents the mean of 3 replicates.

### Validation of protocol

To ensure reproducibility, we provide the following validation data from our internal experiments using the MDA-MB-231 cell line:

- Spheroid formation & consistency: Using the agarose-coated plates described, we consistently formed compact, spherical aggregates within 24 hours. Spheroids remained stable and increased in size progressively over a 96-hour culture period.
- Quantitative analysis: Spheroid size (circumference) was measured daily (n = 12 spheroids per seeding density) using ImageJ software. Data are presented as mean ± standard deviation (SD) of 3 biological replicates. Statistical analysis was performed using GraphPad Prism 9, with ordinary one-way ANOVA applied to compare size across different time points and seeding densities (3×10^3^ to 30×10^3^ cells/well).
- Optimal seeding density: A seeding density of 1.0 x 10^4^ cells/well was identified as optimal, reliably producing a single, central spheroid per well with a uniform spherical morphology suitable for subsequent drug treatment assays.
- Controls: Control wells containing culture medium only were included in every plate to confirm the absence of contamination.

This protocol has been used and validated in the following research article(s):

- **Cervantes-Rivera, R. *et al***. (Manuscript in preparation). Evaluating the antitumor potential of *Persea americana* var. drymifolia seed lipid extract using MDA-MB-231 cell Cultures.

## Acknowledgment

We gratefully acknowledge funding from Coordinación de la Investigación Científica de la UMSNH (Proyecto 17644). We are grateful for stipend support from Secretaría de Ciencia, Humanidades, Tecnología e Innovación (SECIHTI).

